# CRISPR-Cas12a bends DNA to destabilize base pairs during target interrogation

**DOI:** 10.1101/2024.07.31.606079

**Authors:** Katarzyna M. Soczek, Joshua C. Cofsky, Owen T. Tuck, Honglue Shi, Jennifer A. Doudna

## Abstract

RNA-guided endonucleases are involved in processes ranging from adaptive immunity to site-specific transposition and have revolutionized genome editing. CRISPR-Cas9, -Cas12 and related proteins use guide RNAs to recognize ∼20-nucleotide target sites within genomic DNA by mechanisms that are not yet fully understood. We used structural and biochemical methods to assess early steps in DNA recognition by Cas12a protein-guide RNA complexes. We show here that Cas12a initiates DNA target recognition by bending DNA to induce transient nucleotide flipping that exposes nucleobases for DNA-RNA hybridization. Cryo-EM structural analysis of a trapped Cas12a-RNA-DNA surveillance complex and fluorescence-based conformational probing show that Cas12a-induced DNA helix destabilization enables target discovery and engagement. This mechanism of initial DNA interrogation resembles that of CRISPR-Cas9 despite distinct evolutionary origins and different RNA-DNA hybridization directionality of these enzyme families. Our findings support a model in which RNA-mediated DNA engineering begins with local helix distortion by transient CRISPR-Cas protein binding.

## INTRODUCTION

CRISPR-Cas12a uses guide RNAs to identify complementary ∼20-nucleotide sequences in genomic DNA to facilitate binding and double-strand cleavage (1). To engage DNA, Cas12a probes nucleotides immediately adjacent to 5’-TTTV-3’ protospacer adjacent motifs (PAMs) to test for guide RNA hybridization (1–3). When a sequence match is found, RNA-DNA binding forms an R-loop structure, enabling the RuvC nuclease domain to make staggered DNA cuts (1, 2). In bacteria, double-stranded DNA breaks often induce DNA degradation, whereas in eukaryotic cells such breaks can trigger genome editing due to DNA repair (4, 5).

The precise mechanism by which Cas12a locates target sequences amidst the vast excess of non-target sites in a typical genome remains unclear. Structural studies of both Cas9 and Cas12a revealed conformational changes that accompany R-loop formation but did not identify the initial steps of DNA engagement (6, 7). Structural and biochemical studies of Cas9-guide RNA in complex with PAM-containing but otherwise non-target DNA provided evidence for DNA bending causing transient melting of the DNA sequence immediately adjacent to PAMs, enabling initial RNA-guided sequence interrogation (8). Because Cas9 and Cas12 evolved independently from different ancestral proteins (9, 10), it has been unclear whether Cas12a functions by a similar mechanism (Fig. 1A).

**Figure 1.**
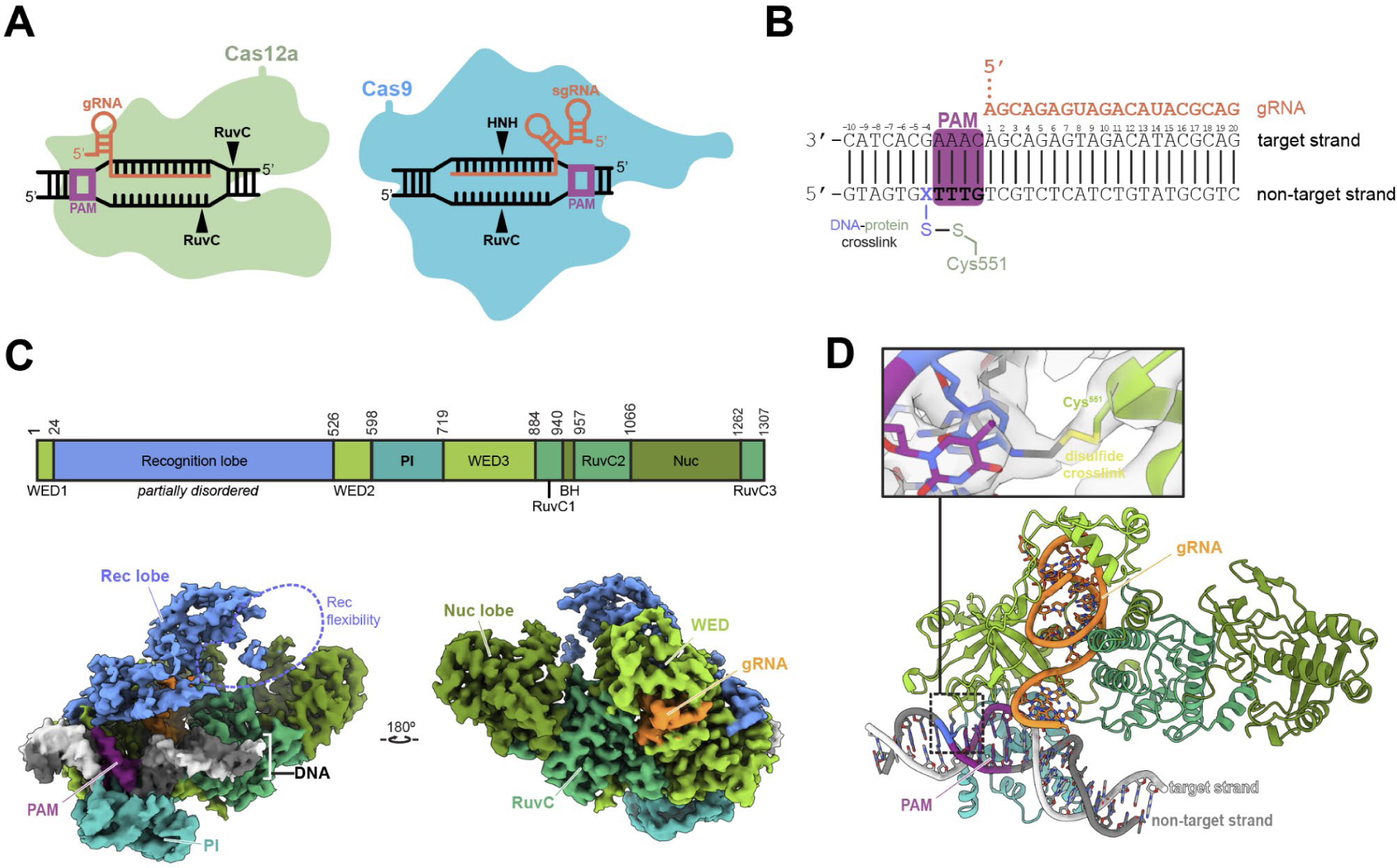
Disulfide crosslinking captures transient protein - DNA interactions. **A.** Schematic representation of Cas12 (left) and Cas9 (right) R-loop complexes highlighting the differences between both enzymes. **B**. DNA and guide RNA spacer sequences used in the study. The PAM sequence is highlighted in green; X shows the position of the cystamine modified base. **C.** AsCas12a domain organization and cryoEM density of sharpened map of structure 2. **D.** Model of structure 2 with detail centered on the disulfide bond crosslink between Cas12 and DNA. The Rec lobe is hidden for clarity.

We used cryo-electron microscopy (cryo-EM) to determine structures of a complex between *Acidaminococcus sp.* Cas12a-guide RNA and a PAM-containing non target DNA molecule. Using site-specific cross-linking to trap the otherwise transient encounter between the Cas12a ribonucleoprotein (RNP) and the DNA, we observed three classes of DNA conformations with different degrees of bending relative to a standard B-form helix. In the most distorted class, a PAM-proximal target strand nucleotide is unstacked and dynamic, an observation supported by fluorescence-based measurements of DNA conformation in solution. These data suggest that DNA bending induces base-flipping to enable Cas12a RNA-guided target recognition, a mechanism analogous to that of Cas9 despite independent evolutionary origins (8). Our findings help explain how Cas12a identifies target sites within genomes, a process that influences both the rate and outcomes of Cas12a-mediated genome editing.

## MATERIALS AND METHODS

### Protein expression and purification

Protein expression and purification were conducted as described in Cofsky et al., 2020. Briefly, BL21 Star (DE3) *E. coli* (Invitrogen) transformed with pJCC_099 was grown in Terrific Broth (TB) and induced at mid-log stage growth with 0.5 mM IPTG overnight at 16°C. Cell pellets were resuspended in lysis buffer (50 mM HEPES (pH 7.5), 500 mM NaCl, 1 mM TCEP, 0.5 mM PMSF, 10 tablets/L cOmplete EDTA-free protease inhibitor cocktail (Roche), and 0.25 mg/mL chicken egg white lysozyme (Sigma-Aldrich)) and sonicated. Lysate was centrifuged, and supernatant was loaded onto Ni-NTA Superflow resin (Qiagen). Resin was washed with wash buffer (50 mM HEPES (pH 7.5), 500 mM NaCl, 1 mM TCEP, 5% glycerol, 20 mM imidazole), and protein was eluted with elution buffer (50 mM HEPES (pH 7.5), 500 mM NaCl, 1 mM TCEP, 5% glycerol, 300 mM imidazole). TEV protease was added to the eluate, which was then dialyzed overnight against dialysis buffer (50 mM HEPES (pH 7.5), 250 mM NaCl, 1 mM TCEP, 5% glycerol). The sample was loaded onto MBPTrap HP and HiTrap Heparin HP columns (Cytiva) connected in series. After removing the MBPTrap column, the Heparin column was washed with ion exchange buffer A (50 mM HEPES (pH 7.5), 250 mM KCl, 1 mM TCEP, 5% glycerol) and eluted with a gradient of ion exchange buffer B (50 mM HEPES (pH 7.5), 1 M KCl, 1 mM TCEP, 5% glycerol). Peak fractions were concentrated and run in gel filtration buffer (20 mM HEPES (pH 7.5), 200 mM KCl, 1 mM DTT, 5% glycerol) over a HiLoad 16/600 Superdex 200 pg column (Cytiva). Peak elution fractions were pooled and concentrated to a final concentration of 80 µM.

### *In vitro* transcription of guide RNA

DNA templates for *in vitro* transcription were assembled by PCR from overlapping DNA oligonucleotides (IDT) (5’-GTCGAAATTAATACGACTCACTATAGG-3′, 5’-AATACGACTCACTATAGGTTTAATTTCTACTCTTGTAGAT-3′, 5’ – CTGCGTATGTCTACTCTGCTATCTACAAGAGTAGAAAT-3′). The transcription reaction was assembled in transcription buffer (40 mM Tris-Cl (pH 7.9 at 25°C), 25 mM MgCl_2_, 10 mM dithiothreitol, 0.01% (v/v) Triton X-100, 2 mM spermidine, 5 mM each NTP, 100 µg/mL T7 RNA polymerase) and allowed to proceed at 37°C for 2.5 hr. DNase I was added, and the sample was incubated for an additional 30 min at 37°C. RNA was then purified by denaturing PAGE (10% acrylamide:bis-acrylamide 29:1, 7 M urea, 0.5X TBE), ethanol-precipitated, and resuspended in RNA storage buffer (0.1 mM EDTA, 2 mM sodium citrate, pH 6.4).

### DNA oligonucleotide preparation

The cystamine-functionalized DNA oligonucleotide (5’-GTAGTGXTTTGTCGTCTCATCTGTATGCGTC, where X denotes the N4-cystamine-2’-deoxycytidine) was synthesized by TriLink Biotechnologies. All other oligonucleotides were obtained from Integrated DNA Technologies. DNA oligonucleotides were purified in-house on urea-PAGE gels. DNA duplexes were annealed in water by heating to 95°C for 2 min and then cooling to 25°C over a 40 min period.

### DNA cleavage assay

Reactions were conducted in DNA cleavage buffer (20 mM Tris pH 7.9, 150 mM KCl, 5 mM MgCl_2_, 5 mM TCEP). Reactions were initiated by adding DNA (non-target strand 5’-GTCATAATGATTTTATCTTCTGGATTGTTGTAAGCAGCATTTGAGCAAAAATCTGTTGC, target strand 5’-GCAACAGATTTTTGCTCAAATGCTGCTTACAACAATCCAGAAGATAAAATCATTATGAC) to AsCas12a:guide RNA complex, yielding final concentrations: 100 nM AsCas12a,120 nM guide RNA, 1 nM radiolabeled DNA duplex. Reactions were quenched by addition of one volume of 2x quench buffer (94% formamide, 30 mM EDTA, 0.025% w/v bromophenol blue) and analyzed by denaturing PAGE (15% acrylamide bis-acrylamide 29:1, 7 M urea, 0.5X TBE) and phosphorimaging.

### Oligonucleotide radiolabeling

T4 polynucleotide kinase (New England Biolabs) at 0.2 U/µL (manufacturer’s units) was mixed with 1X T4 PNK buffer (New England Biolabs), 400 nM DNA oligonucleotide, and 200 nM [γ-^32^P]-ATP (PerkinElmer) and incubated for 30 min at 37°C, then 20 min at 65°C. Radiolabeled oligo was then buffer exchanged into water using a Microspin G-25 spin column (GE Healthcare).

### Complex preparation

For crosslinking optimization, 6 µM functionalized DNA duplex, 5 µM RNA, and 4 µM AsCas12a were combined in cross-linking buffer (50 mM Tris pH 7.4, 150 mM NaCl, 5 mM MgCl_2_, 5% glycerol, 100 µM DTT). The reaction was incubated at 25°C for 12 hr and then quenched by addition of S-methyl methanethiolsulfonate to a final concentration of 20 mM. Reactions were then denatured by adding non-reducing SDS loading solution and heating for 5 min at 90°C.

For cryo-EM sample preparation, reactions were prepared as above but without quenching. After the 12-hour incubation, the sample was injected onto a Superdex 200 Increase 10/300 GL column equilibrated with complex buffer (20 mM Tris pH 7.5, 200 mM KCl, 5 mM MgCl_2_, 0.25% glycerol and 100 µM DTT). Peak fractions were collected, concentrated, and frozen at -80°C for storage.

For 2-aminopurine assays, reactions comprising 1.3 µM Cas12a, 1.7 µM RNA and 1 µM DNA in the buffer specified above were incubated overnight at 25°C to facilitate crosslinking.

### EM grid preparation and data collection

Sample aliquots were thawed and diluted into complex buffer to a concentration of 3 µM. Grids (1.2-µm/1.3-µm 400 mesh C-flat grids, Electron Microscopy Sciences #CF413-50) were glow discharged (PELCO easiGlow) for 15 s at 25 mA. 3.6 µL of sample was applied to a grid in an FEI Vitrobot Mark IV operated at 8°C and 100% humidity. Excess sample was blotted for 3 s with blot force 6 before plunge freezing. Micrographs were collected on a Talos Arctica transmission electron microscope operated at 200 kV and ×36,000 magnification (1.115 Å per pixel), with −0.8 to −2 µm defocus, using the super-resolution camera setting (0.5575 Å per pixel) on a Gatan K3 Direct Electron Detector. 1,684 movies were collected using beam shift in SerialEM v.3.8.7 software.

### Cryo-EM data processing

Initial data processing was conducted using cryoSPARC v.4 (Structura Biotechnology Inc.) (11). After motion correction (Patch Motion) and CTF estimation (Patch CTF), 1,364 micrographs were curated for further processing. Initial particle picking was done using blob picker on a small subset of micrographs to generate templates that were used (Template Picker) to pick 1,929,565 particles from all micrographs. After two rounds of 2D classification, 459,736 particles remained. Particles were re-extracted with re-centering and used for *ab initio* reconstruction into 5 classes. Only one class corresponded to the expected complex (137,851 particles). Particles from this class were again re-extracted with re-centering, and subjected to non-uniform refinement, which provided a map with global resolution of 3 Å. In the sharpened map the DNA density was absent. Particles from the non-uniform refinement were transferred to RELION v. 5.0-beta (12) for DNA-focused refinement. After 3D refinement in RELION, signal from the protein was subtracted, the particles were re-centered on DNA, and five DNA volume classes were reconstructed. Three classes, with 20,031, 27,147 and 27,031 particles respectively, were chosen for final reconstruction. Particle subtraction was reversed to yield reconstructions of the full particles. Reconstructions were refined with masking, solvent-flattened FSC and Blush regularization. Refined maps were sharpened (B-factor -30), reaching resolutions 3.6 Å for structure 1, 3.4 Å for structure 2 and 3.5 Å for structure 3. These maps were used for model building and refinement.

### Model building

The AsCas12a crystal structure (PDB 5B43) was used as an initial model. In Coot (v. 0.9.8.93 EL) (13) each protein domain was fit separately into the density with manual adjustments. Both sharpened and unsharpened maps were used in this process. The protein model was iteratively refined in Phenix (v. 1.19.2-4158) (14–16) using phenix.real_space_refine (minimization_global, local_grid_search, adp) with secondary structural restraints and manual geometry adjustments in Coot. Regions where neither the sharpened nor unsharpened map provided sufficient density remained unmodeled. Overall, the REC domain had the weakest density in the structures. The RNA sequence was adjusted to reflect our construct sequence. DNA was created as a linear B-form helix in Coot. PHENIX (16) simulated_annealing with base pair and base stacking restraints was used to fit the DNA into the density. In one of the structures, the base near the PAM sequence was modeled as unstacked and rotated away from the helix axis. UCSF Chimera X ISOLDE (17) was used to improve the model of the DNA, as well as the PI and REC domains of Cas12a. Planarity was enforced on nucleotides within the DNA duplex in the protospacer region (5 base pairs; chain C/D bases 14-18) for all structures. The thioalkane linker was added as an ethanethiol ligand from PHENIX’s eLBOW module (18). Sulfur-sulfur and carbon-nitrogen bonds were restrained as described (8).

### 2-Aminopurine assay to detect unstacked nucleotides

Target strand oligonucleotides including a 2-aminopurine base in position 1 (5’-GACGCATACAGATGAGACG**A**<u>CAAA</u>GCACTAC-3′ – bold font shows the modification insertion, underlined nucleotides are the PAM), position 2 (5’-GACGCATACAGATGAGAC**G**A<u>CAAA</u>GCACTAC-3′), position 4 (5’-GACGCATACAGATGAG**A**CGA<u>CAAA</u>GCACTAC-3′) and position 13 (5’-GACGCAT**A**CAGATGAGACGA<u>CAAA</u>GCACTAC-3′) downstream of the PAM sequence in the protospacer region, were obtained from IDT. Reactions (in triplicate, 25°C) were assembled and measured every 75 s over 2 hours using a Biotek Cytation 5 imaging reader with excitation wavelength 320 nm and emission wavelength 370 nm. Reading at 75 sec point was used for analysis. Reading of triplicates were averaged together. All measurements were normalized to the signal from double stranded DNA alone and measurement error was propagated using the following formula: 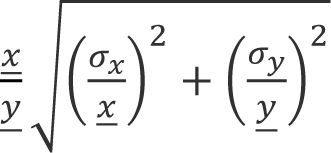 where <u>*x*</u> is the average of Cas12a-RNA-DNA or Cas12a-DNA signal; <u>*y*</u> is the average of DNA only signal; σ_*x*_ is a standard deviation for Cas12a-RNA-DNA or Cas12a-DNA signal; σ_*y*_ is a standard deviation for DNA only signal.

### DNA bend and twist calculations

We quantified the local DNA bending and unwinding simultaneously, using an established inter-helical Euler angle approach (19, 20). This method measures the bending magnitude (β_h, 0° ≤ β_h ≤ 180°), bending direction (γ_h, -180° ≤ γ_h ≤ 180°), and helical twist (ζ_h, -180° ≤ ζ_h ≤ 180°) between two helices (H1 and H2) across a junction (J) containing one or multiple base pairs. We defined the PAM sequences as H1, the spacer sequences two base pairs away from PAM as H2 and the two base pairs immediately adjacent to PAM in the spacer as J. Two idealized B-form DNA helices, each comprising 3 base pairs, were constructed using the 3DNA software (21) and superimposed onto H1 and H2, respectively. This alignment enabled us to determine the relative orientation of H1 to H2, quantified by the parameters β_h, γ_h, and ζ_h. The underwinding angle of the helix was calculated by N x 36° -ζ_h, where N is the number of base pairs in the J. Using this method, bending angle for Cas9 interrogation complex with bent DNA and closed protein conformation had a bend angle ∼69° and unwinding of ∼66°, while the linear DNA conformation in an open protein conformation had a bend angle of ∼38° and unwinding of ∼16°.

### Figure preparation

The comparison of PI domains was conducted after structure alignment with SSM superpose in Coot using protein chain. For figure visualization models were aligned with Chimera X (1.7.1) (22) Matchmaker using protein chain in the reference for alignment. DNA-protein contacts were listed with Chimera X “contacts” command. Figures were prepared with Chimera X (1.7.1). Chromatogram of complex purification and 2-AP assay results were plotted using GraphPad Prism 8.

## RESULTS

### Cryo-EM structures of a crosslink-stabilized Cas12a:guide RNA:non target DNA complex

To investigate target search by a Cas12a-guide RNA complex, we designed DNA substrates bearing a 5’-TTTG-3’ PAM sequence but lacking any base pairing complementarity with the guide RNA (Fig. 1B). We captured transient Cas12a RNP association with this substrate by introducing a disulfide crosslink between mutated amino acid N551C in Cas12a and an N4-cystamine-cyt(7) DNA modification. Control experiments confirmed that N551C mutation in Cas12a did not disrupt its ability to cleave target DNA (Fig. S1A). Formation of the disulfide crosslink between Cas12a and the modified DNA was confirmed by denaturing, non-reducing SDS-PAGE analysis (Fig. S1B).

Cryo-EM analysis of the cross-linked sample revealed three distinct molecular structures (Fig. S1C, S2, Table 1). Conformations of the protein in each model are similar to each other and resemble Cas12a-guide RNA binary complexes (PDB ID 5NG6, 5ID6). In all structures, the PAM sits in the Cas12a PAM-binding pocket (Fig. 1C). EM maps show clear density corresponding to the crosslinking disulfide bridge between the protein and the DNA (Fig. 1D).

**Table 1.** CryoEM data collection, refinement and validation statistics.

| Data collection |  |  |  |
| --- | --- | --- | --- |
| sample | Structure 1<br>PDB 9CJH<br>EMD-45631 | Structure 2<br>PDB 9CJI<br>EMD-45632 | Structure 3<br>PDB 9CJJ<br>EMD-45633 |
| Magnification | 36,000 | 36,000 | 36,000 |
| Voltage (kV) | 200 | 200 | 200 |
| Electron dose (e-/Å<sup>2</sup>) | 50 | 50 | 50 |
| Defocus range (μm) | -0.8 to 2 | -0.8 to 2 | -0.8 to 2 |
| Pixel size (Å) | 1.115 | 1.115 | 1.115 |
| **3D reconstruction** |  |  |  |
| Raw images | 1364 | 1364 | 1364 |
| Initial particles | 1,929,565 | 1,929,565 | 1,929,565 |
| Final particles | 20,031 | 27,147 | 27,031 |
| Map resolution (Å) | 3.6 | 3.4 | 3.5 |
| FSC threshold | 0.143 | 0.143 | 0.143 |
| **Model refinement** |  |  |  |
| Initial model used | Structure 2 | 5B43 | Structure 2 |
| Model resolution |  |  |  |
| FSC threshold |  |  |  |
| <u>Model composition</u> |  |  |  |
| Nonhydrogen atoms | 10949 | 10908 | 10744 |
| Protein residues | 1158 | 1158 | 1158 |
| Nucleotide | 69 | 67 | 59 |
| Ligands | 1 | 1 | 1 |
| <u>B factors (mean, Å<sup>2</sup>)</u> |  |  |  |
| Protein | 75.36 | 69.75 | 74.06 |
| Nucleotide | 137.39 | 101.99 | 100.63 |
| Ligand | 98.79 | 76.23 | 84.16 |
| <u>R.m.s deviations</u> |  |  |  |
| Bond length (Å) | 0.003 | 0.003 | 0.003 |
| Bond angles (°) | 0.506 | 0.520 | 0.525 |
| **Validation** |  |  |  |
| MolProbity score | 1.63 | 1.45 | 1.58 |
| Clash score | 8.98 | 8.1 | 9.21 |
| Rotamer outlier (%) | 0 | 0 | 0 |
| <u>Ramachandran statistics (%)</u>- |  |  |  |
| Favored | 97.19 | 97.98 | 97.54 |
| Allowed | 2.81 | 2.02 | 2.46 |
| Outlier | 0 | 0 | 0 |
| Rama-Z score, whole<br>(r.m.s Rama-Z) | 1.8<br>0.26 | 1.83<br>0.26 | 1.72<br>0.27 |
| Map CC (box) | 0.72 | 0.74 | 0.73 |
| Map CC (mask) | 0.76 | 0.78 | 0.77 |

Consistent with the flexibility of the recognition (REC) lobe in both Cas12a and Cas9 noted in prior studies (6–8, 23, 24), we observed poor EM density for the REC domain in the structures determined here.

### Movement of PAM interaction domain induces DNA bending

We analyzed Cas12a-DNA interactions in our cryoEM structures to assess their similarity to prior Cas12a complexes containing target-engaged Cas12a RNPs (2, 7, 25). In all structures, the DNA sits in a positively charged groove of PI and WED domains. In structures 2 and 3, Pro599, Met604 and Lys607 of the PAM-interacting (PI) domain and Lys548 of the Wedge (WED-II) domain form contacts with the PAM directly adjacent to the crosslink position (Fig. 2A) (2, 3). In structure 1, key interactions between Lys548 and the PAM are absent. N7 of adenosine (position -2) on the target strand is 4.46 Å away from Lys548 side chain, placing it out of range for hydrogen bonding. The majority of protein-DNA contacts surround PAM interaction residues and form non-specific interactions with the DNA backbone. This is consistent with previously published molecular structures of the ternary complex of AsCas12a determined by X-ray crystallography (PDB ID 5B43) and cryoEM (PDB ID 8SFH).

**Figure 2.**
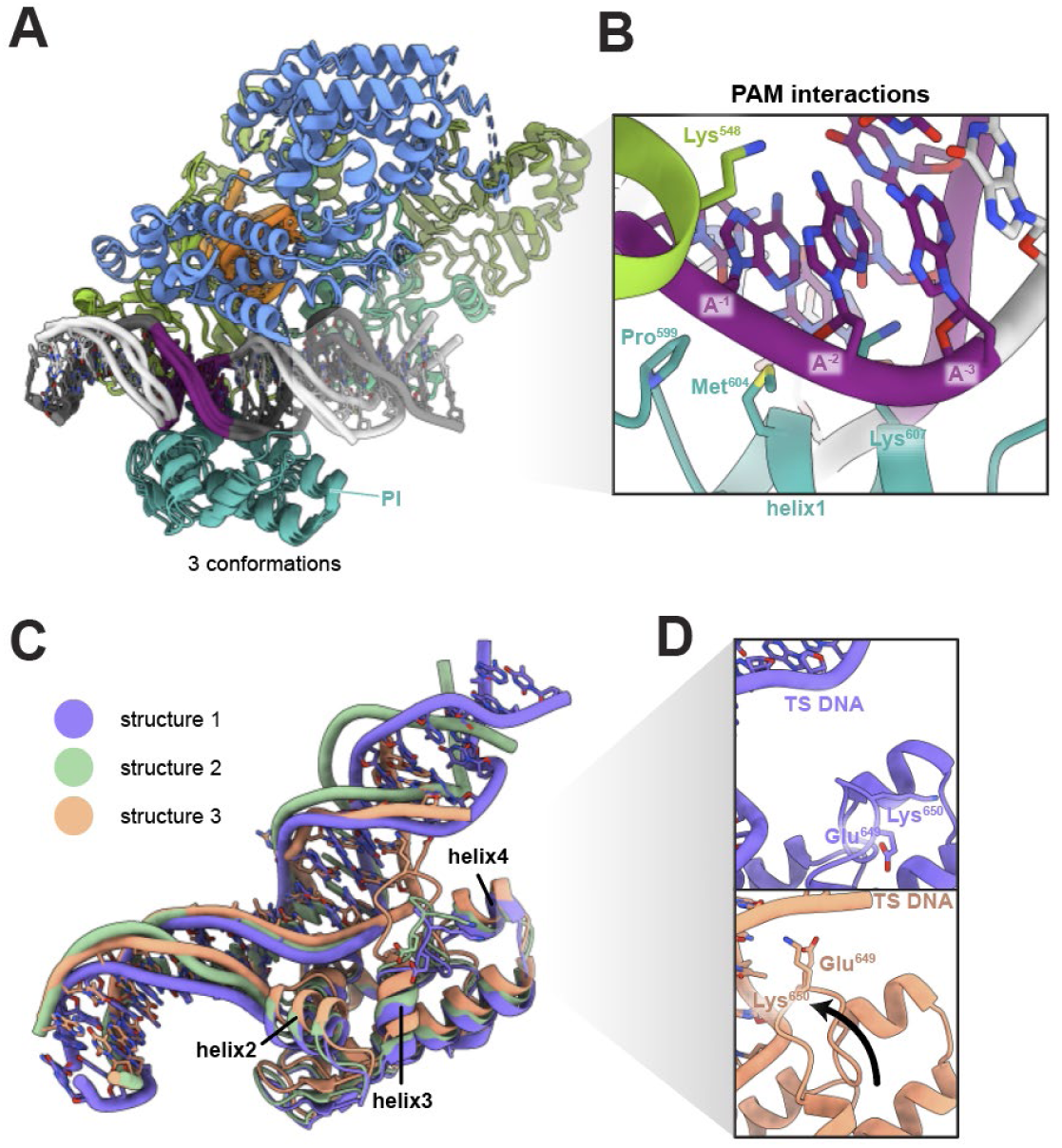
Cas12a - DNA interactions and PI domain movements. **A.** Global protein alignment of all structures. **B**. Detail of PAM recognition interactions. View from the target strand side. **C.** Close-up of the PI domain and DNA conformational rearrangements. **D.** Comparison of loop between helix 3 and 4 in structure 1 (top) and structure 3 (bottom).

The C-alpha alignment of all three molecular models shows the PI domain of structure 2 shifts towards the DNA by less than 1 Å globally. Helix 2 moves by ∼1.3 Å relative to in structure 1. This shift is accompanied by an intermediate DNA conformation with a more pronounced bending and underwinding (45° bending and 19° underwinding versus 32° bending and 3° underwinding) than in structure 1 (Fig. 2C, 3A). Structure 3 displays the largest relative rearrangement of the PI domain. Relative to structure 1 the entire PI domain of the structure 3 folds towards the DNA. The largest rearrangement is visible in helices 1-4. Helix 1, which hydrogen bonds with the PAM, moves towards DNA by ∼1.5 Å. Helix 3 moves towards the DNA by ∼3 Å, with concomitant motion of helix 2 by ∼2.2 Å (Fig. 2C). The result is a large bending and underwinding of the DNA of 68° and 41°, respectively (Fig. 2C, 3A). In addition, a loop between helix 3 and 4 in the structure 3 reaches towards the PAM-distal fragment of the DNA and may form transient interactions with the DNA backbone (Fig. 2C,D). These results show that movements of the PI domain correlate with DNA bending and underwinding.

### Cas12a bends DNA to induce helical distortion and base flipping for DNA interrogation

We quantified the relative distortion of DNA in crosslinked structures (Fig. 3A). DNA in the candidate protospacer region is progressively bend and underwound from structures 1 to 3. In structure 3, which displays a DNA bend of 63° and is underwound by 41°, we observe base unstacking and flipping (Fig. 3B-E). The nucleobase in position 2 downstream of the PAM within the candidate protospacer region is rotated out of its normal DNA base-pairing position (Fig. 3A, E; Fig. S3). The corresponding nucleobase on the opposite strand (NTS) is also slightly unstacked. The absence of the flipped base in structures 1 and 2 suggest base flipping arises from protein induced DNA bending.

**Figure 3.**
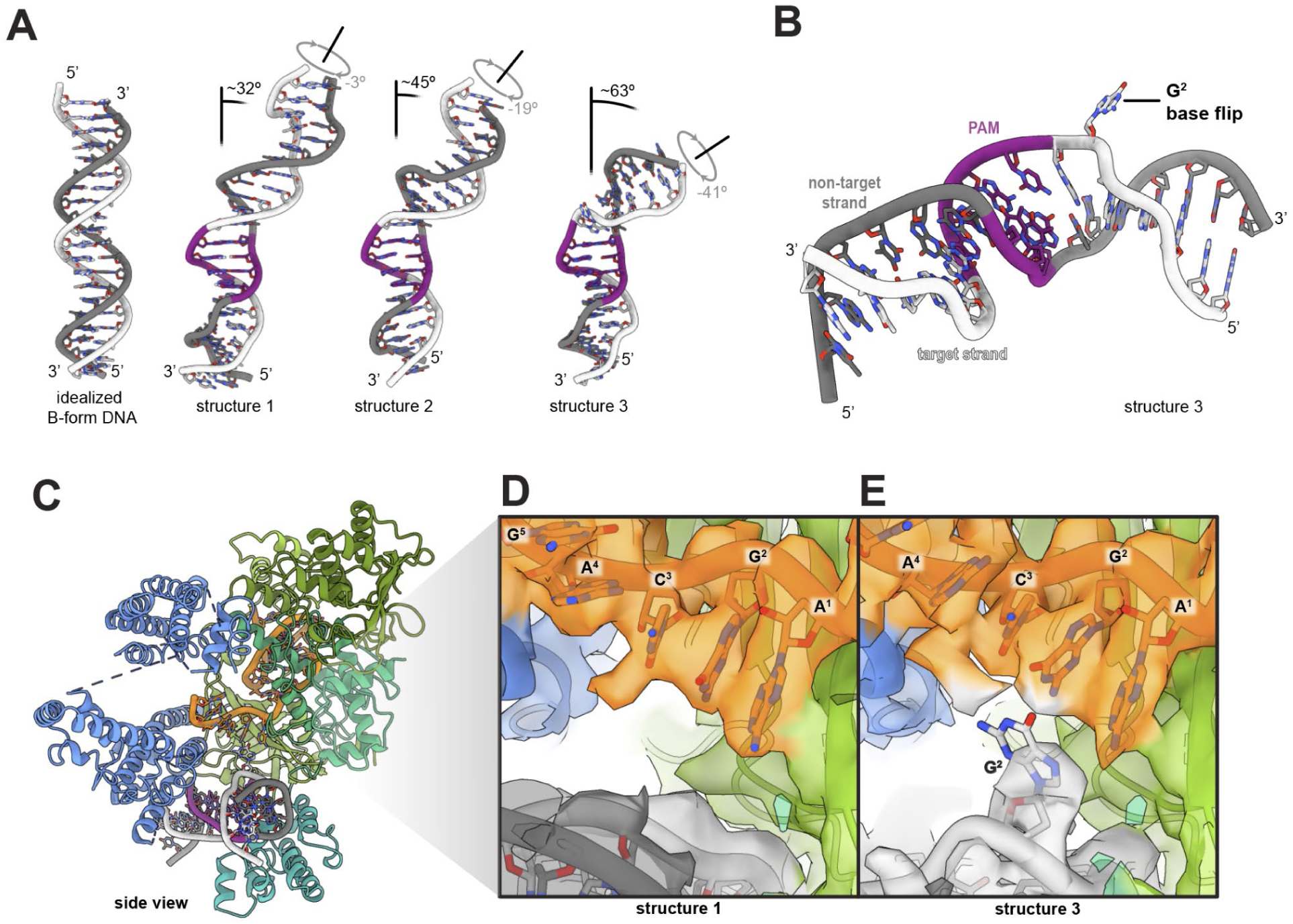
DNA bending and base flipping. **A.** Comparison of B-form DNA with all DNA bent and underwound conformations. **B.** View of Structure 3 DNA showing flipped base. **C.** Side view of structure 3. **D.** Close-up of the density in proximity to the RNA seed sequence in structure 1. **E.** Close-up of the density in proximity to the RNA seed sequence in structure 3.

To investigate whether DNA base flipping occurs during Cas12a DNA interrogation, we employed a fluorescence-based biochemical assay. We designed DNA substrates containing single 2-aminopurine (2-AP) nucleotides at different positions relative to the PAM (Fig. 4A). This assay monitors the increase in fluorescence that occurs when 2-AP moves from a stacked, base-paired environment to an unstacked, single-stranded environment (Fig. 4A). In control reactions containing Cas12a crosslinked to DNA in the absence of guide RNA, we did not observe an increase in fluorescence relative to the 2-AP-containing double stranded DNA alone. We next conducted fluorescence detection assays using Cas12a RNPs crosslinked to a non-targeting sequence. We observed a pronounced increase in fluorescence for substrates with 2-AP at positions 1 or 2 downstream of the PAM. Only a small increase in fluorescence was observed for substrates with 2-AP at positions 4 and 13 (Fig. 4B). These data are consistent with our structural data suggesting Cas12a flips DNA bases independent of spacer complementarity during target interrogation. A recent study of the Cas9 DNA search mechanism provided evidence for solvent exposure of bases 1 and 2 adjacent to the PAM (8). Similarity between Cas9 and Cas12a suggests that base flipping induced by DNA bending is a common interrogation strategy for Class 2 CRISPR-Cas effectors.

**Figure 4.**
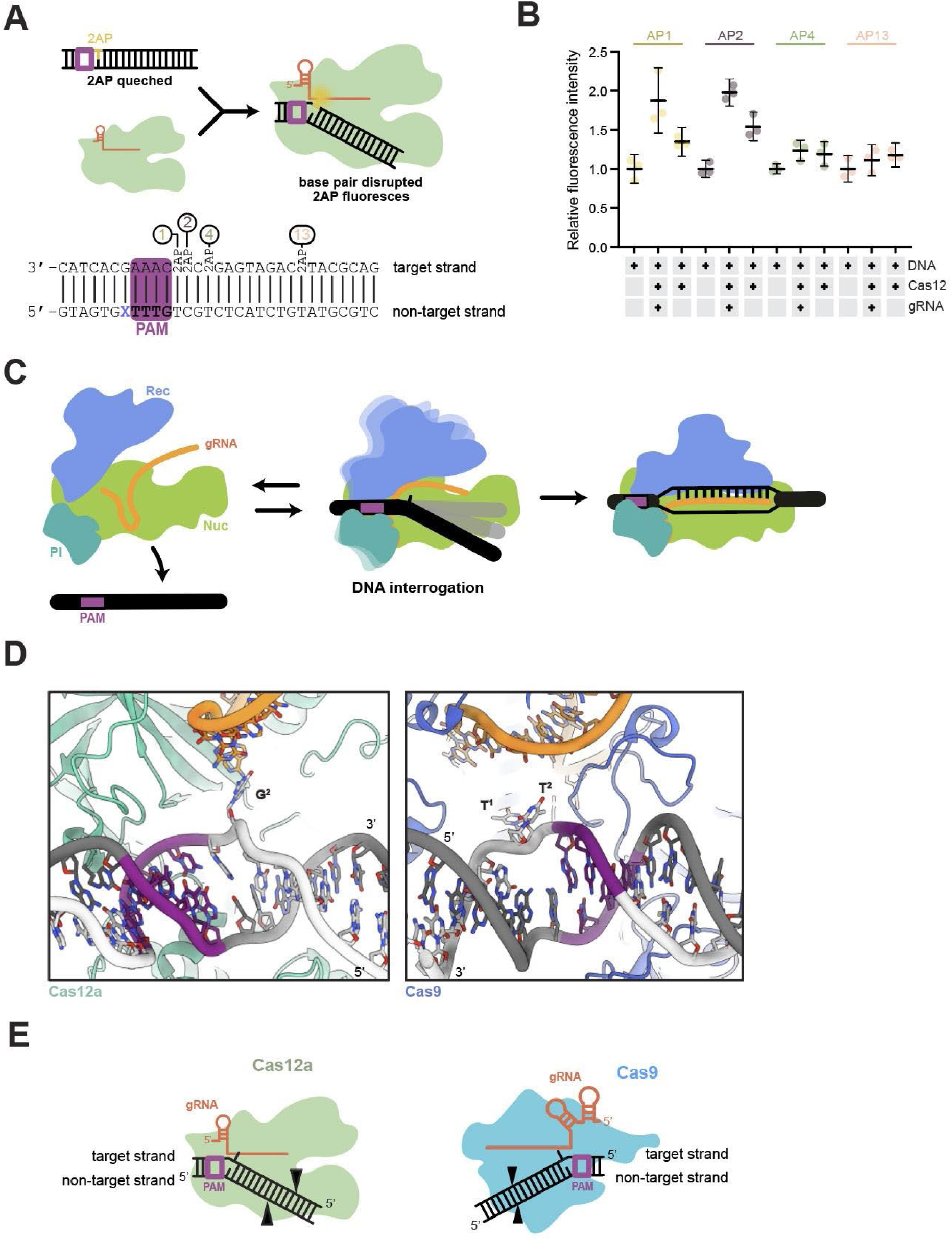
Cas12a interrogation mechanism. **A.** Schematic representation of the 2-AP assay experimental design and expected outcomes. DNA sequence tested with positions of 2-AP insertion in all probes. **B.** Graph showing triplicate fluorescence measurement for double stranded DNA, Cas12-DNA-RNA complex and Cas12-DNA control. The average of the triplicate signal normalized to the average of the double stranded DNA signal are plotted with error bars as described in methods. **C.** Bent DNA state symbolizes the dynamic conformation solved in this study. **D.** Close-up of the PAM proximal region of Cas12a structure 3 (this study, left) and Cas9 interrogation complex (PDB:7S36, right). **E.** Schematic representation of Cas12a (left) and Cas9 (right) in DNA bending and base flipping state.

## DISCUSSION

In this study, we examined the early stage of DNA interrogation by Cas12a. We trapped a normally transient DNA binding interaction to unravel the mechanism of initial DNA recognition before R-loop formation. Cryo-EM analysis of a cross-linked Cas12a-guide RNA-DNA complex revealed three distinct structural views representing the initial interaction between Cas12a and DNA. The DNA in each structure binds similarly to the PAM interacting (PI) domain, but differs in the degree of distortion from the canonical B-form helix. DNA bending appears to be induced by subtle conformational changes in the Cas12a PI domain. We observed the largest PI domain repositioning in structure 3, where helices and a bridging loop shift towards DNA. Protein motions accompany DNA bending and underwinding which culminates in a base flipping of the second nucleobase of the candidate protospacer region. The flipped base may serve to avoid steric clashes and potentially initiate RNA-guided sequence interrogation. We confirmed conformational changes in a 2-AP assay, where we observed an increase in fluorescence for PAM-proximal, but not PAM-distal, bases.

Analogous structures obtained for the Cas9 interrogation complexes showed that DNA bending correlates with differences in protein conformation (8). In the most distorted DNA structure, DNA was bent and underwound to 68° and 66° respectively, exposing two bases adjacent to the PAM. Although Cas9 introduces more dramatic distortions than Cas12a, both are capable of flipping PAM-adjacent bases through a DNA bending mechanism to interrogate complementarity with the guide RNA. Our results demonstrate that Cas12, like Cas9 (8), induces conformational rearrangements to force DNA into a bent conformation. Ribonucleoprotein-mediated DNA bending causes base flipping and facilitates interrogation of the sequence after PAM binding. Whereas Cas9 interrogates the target sequence through REC and HNH domain movements (6, 8, 27), Cas12a repositions the PI and REC domains.

Cas9 searches for targets using both three-dimensional diffusion (28) and locally facilitated one-dimensional diffusion (29). In contrast, Cas12a searches for targets mainly through one-dimensional diffusion along the DNA and sometimes can even bypass a target site during DNA target searching (30). The cryo-EM model with less bent DNA in this study provides a potential physical explanation for the distinct search mode used by Cas12a. The frequency of Cas12a-induced base flipping may explain how often Cas12a bypasses target sites during 1D diffusion on the DNA.

Together, these data support a model in which Cas12a locates target sites within a genome by engaging PAM sequences and utilizing subtle PI domain conformational changes, along with REC domain movements, to bend and unwind DNA. Introduction of DNA distortion encourages base flipping, making PAM-adjacent nucleobase(s) accessible to potential base pairing with the guide RNA. Without guide complementarity, Cas12a releases the DNA and locates another PAM sequence (Fig. 4C).

Similarities between Cas9 and Cas12a DNA interrogation mechanisms imply convergent evolution of ATP-independent, RNA-mediated DNA interrogation. The evolution of Cas9 and Cas12a from different ancestral proteins (9) implies that transient DNA distortion induced by DNA bending may be a mechanism shared by RNA-guided enzyme families. Local DNA unwinding causes bases adjacent to the PAM to become unstacked and exposed in the direction of the guide RNA. This occurs despite opposite directionalities of the PAM with respect to the candidate protospacer region (Fig. 4D). Both enzyme families use the biophysics of helix destabilization to enable RNA-guided target recognition (Fig 4E). In both their native (31, 32) and biotechnological contexts (33, 34), Cas9 and Cas12 survey numerous DNA sequences for the guide RNA complementarity, limiting efficiency by residency at off-target sites. We propose transient distortion of the PAM proximal region through DNA bending is an efficient biophysical solution to this problem. Cas effectors sample DNA locally before complete R-loop formation, enabling rapid dissociation from non-cognate spacers. These findings help explain why Class II CRISPR Cas effectors function even in eukaryotic genomes many times larger than those of their original prokaryotic hosts. Cas9 and Cas12 enzymes may differ in conformational sampling to tune speed and fidelity in their native contexts (31). These parameters may be utilized to engineer more efficient systems for genome editing efforts.

## DATA AVAILABILITY

Plasmid sequences and structural data will be released upon publication of this manuscript.

## FUNDING

The project was funded by grants from the National Institutes of Health [NIH/NIAID (U19AI171110, U54AI170792, U19AI135990, UH3AI150552, U01AI142817), NIH/NINDS (U19NS132303), NIH/NHLBI (R21HL173710)] and the National Science Foundation [2334028] to J.A.D. . J.A.D. is an investigator of the Howard Hughes Medical Institute (HHMI). K.M.S. was funded in part by the Panattoni Family Foundation. J.C.C. was previously a fellow of the National Science Foundation Graduate Research Fellowship. J.C.C. is currently a fellow of the Helen Hay Whitney Foundation. H.S. is an HHMI Fellow of The Jane Coffin Childs Fund for Medical Research. J.A.D. also acknowledges support from Emerson Collective, Department of Energy [mCafes, DE-AC02-05CH11231, 2553571, B656358], Apple Tree Partners (24180), Lawrence Livermore National Laboratory, UCB-Hampton University Summer Program, Mr. Li Ka Shing, Koret-Berkeley-TAU, and the Innovative Genomics Institute (IGI).

### ACKNOWLEDGEMENTS

We thank members of the Doudna lab and the Innovative Genomics Institute for helpful discussions. We thank Dr. Erin Doherty for assistance with 2-aminopurine assays. We thank N.Moriarty for assistance in modeling thioalkane linker. We thank Jonathan Remis and Daniel Toso for electron microscopy assistance and Abhiram Chintangal for computational support. We thank Prof. Eva Nogales, Alfredo Florez Ariza and all members of the Nogales lab for advice and discussions about cryo-EM data processing. EM data were collected at the Cal-Cryo facility located at UC Berkeley. We also thank Prof. Jamie Cate for critical reading and comments on the manuscript.

## AUTHOR CONTRIBUTIONS

K.M.S., J.C.C. and J.A.D. designed the study and performed the experiments with input and assistance with data analysis from other authors; all authors participated in manuscript and figure preparation and review.

