## Supplementary Data for "CRISPR-Cas12a bends DNA to destabilize base pairs during target interrogation"

[1]Department of Molecular and Cell Biology, University of California, Berkeley; Berkeley, CA, USA; [2]Innovative Genomics Institute; University of California, Berkeley, CA, USA; [3]California Institute for Quantitative Biosciences (QB3), University of California, Berkeley, CA, USA; [4]Department of Chemistry, University of California, Berkeley; Berkeley, CA, USA; [5]Howard Hughes Medical Institute, University of California, Berkeley; Berkeley CA, USA; [6]Gladstone-UCSF Institute of Genomic Immunology; San Francisco, CA, USA; [7]Molecular Biophysics and Integrated Bioimaging Division, Lawrence Berkeley National Laboratory; Berkeley, CA, USA

[#]Present address: Department of Biological Chemistry and Molecular Pharmacology, Harvard Medical School, Boston, Massachusetts 02115, USA

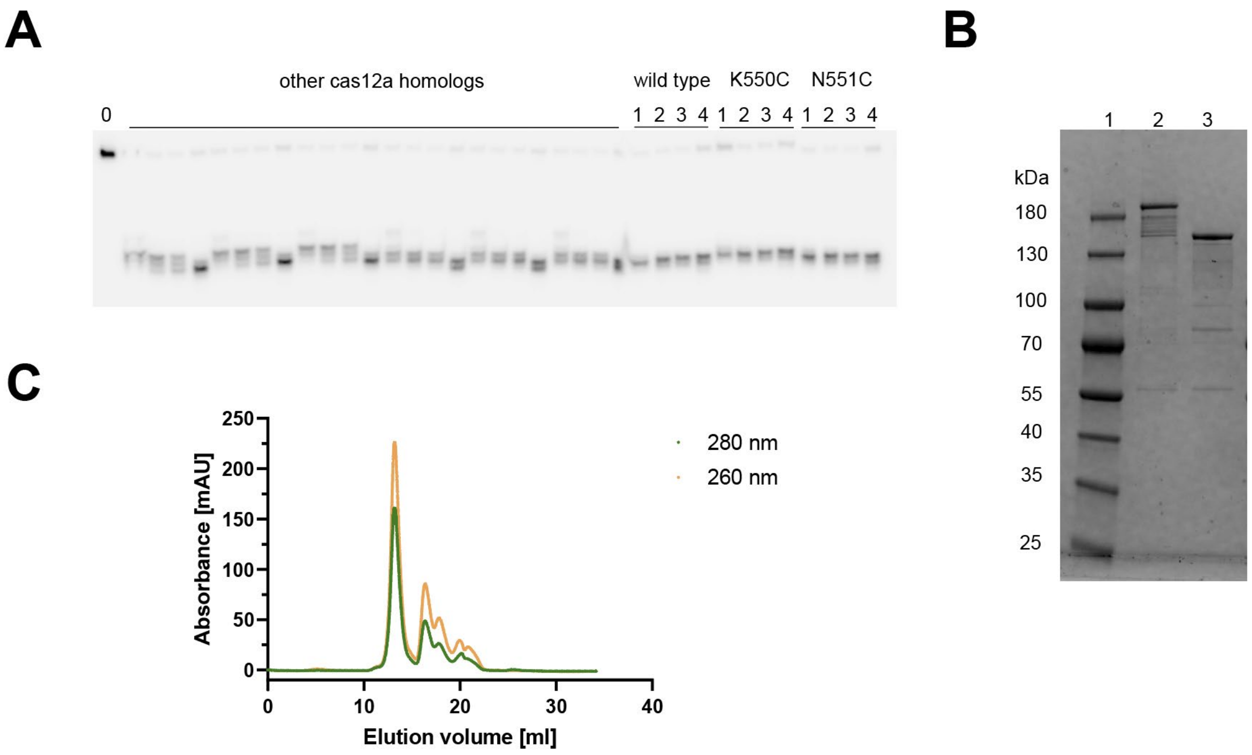


**Supplementary Figure S1. Preparation of the crosslinked complex.**

A. Gel showing cleavage assay of non-target DNA with wild type and mutant AsCas12a. Lane 0 -uncleaved DNA, lanes 1-4 correspond to different reaction incubation times 15 s, 30 s, 1 min, 10 min respectively. **B.** Non-reducing SDS-PAGE gel showing efficiency of Cas12a(N551C)-DNA crosslinking in complex with RNA. Lane 1 protein ladder, lane 2 - crosslinked complex reaction, lane 3 - uncrosslinked protein alone. **C.** Size exclusion complex purification trace prior to cryoEM analysis.

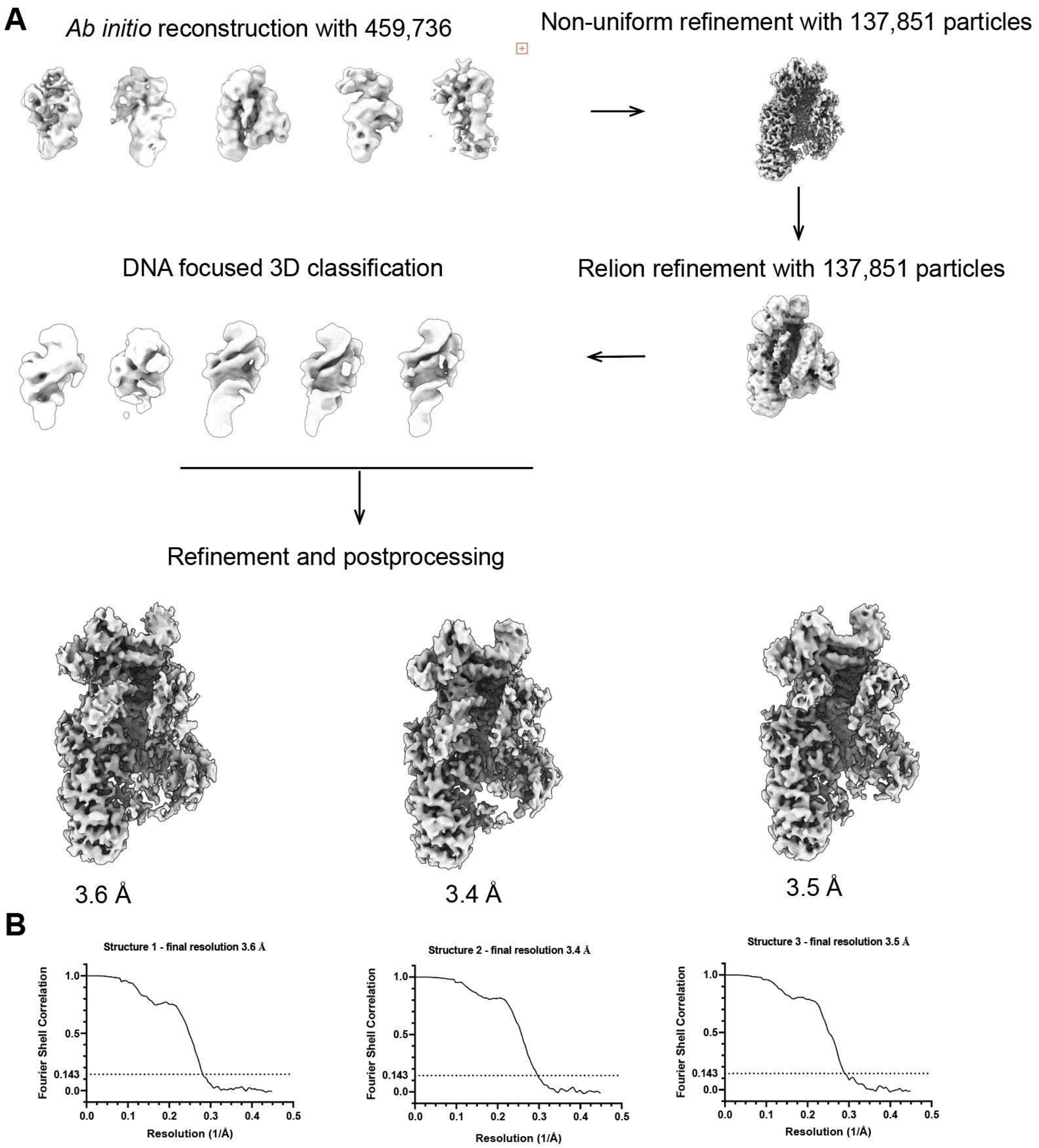


**Supplementary Figure S2. CryoEM data processing.**

**A.** Schematic of particle classification leading to final structures. **B.** FSC curves for each final map.

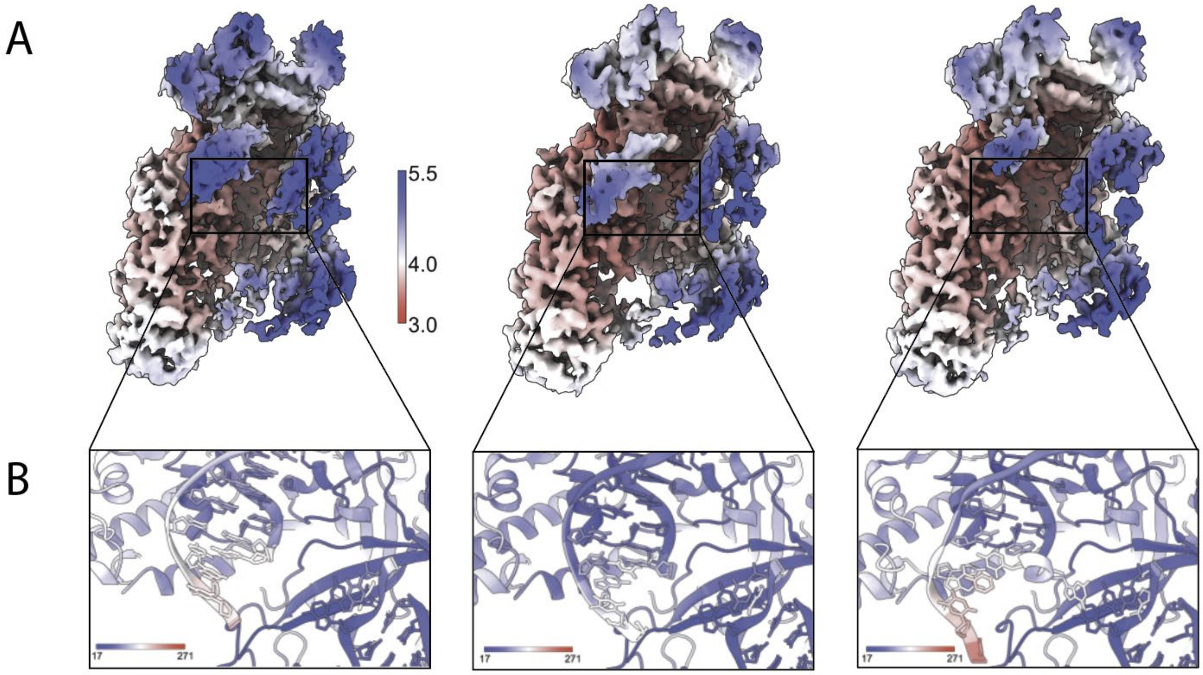


**Supplementary Figure S3. Local structural variability.**

**A.** Maps of all structures in this study colored based on local resolution calculated in RELION 5.

**B.** Close-up on PAM and a region of expected flipped base for all maps, colored based on B-factors.
